# Inheritance of the genome-less apicoplast in the “apicoplast-minus” *Plasmodium falciparum*

**DOI:** 10.1101/2025.03.05.641705

**Authors:** Wei Xu, Ikechukwu Nwankwo, Sean T. Prigge, Hangjun Ke

## Abstract

Most apicomplexan parasites contain a plastid-derived organelle called the apicoplast, which originated through secondary endosymbiosis. As a result of this evolutionary trajectory, the non-photosynthetic apicoplast is surrounded by four membranes and contains many bacterial-like, druggable targets. It is widely accepted that asexual malaria parasites (*Plasmodium falciparum*) can thrive under antibiotic treatment if supplemented with high concentrations of isopentenyl pyrophosphate (IPP, 200 µM) and these IPP-rescued parasites are thought to lack the apicoplast and its 35 kb genome but possess many vesicles. However, our findings challenge this apicoplast-minus concept. In late-stage schizonts, we observed that the apicoplast-derived vesicles nearly colocalize with mitochondria and are properly distributed into merozoites during schizogony, suggesting that they are inherited rather than newly synthesized in each asexual cycle. Further, immuno-electron microscopy (immuno-EM) revealed that the “apicoplast-minus” parasites possess structures surrounded by four membranes, in addition to single-membrane-surrounded entities. The presence of four-membrane-bound structures suggests that the apicoplast has not truly disappeared in the “apicoplast-minus” *P. falciparum* but remains in a distinct, diminished form. We termed this genome-less apicoplast derivative the apicosome, drawing an analogy to the genome-less mitochondrial derivative known as the mitosome. We propose that apicosomes retain essential biochemical and/or structural functions, which act as barriers to the complete loss of apicoplast when the parasites face antibiotic stress and IPP rescue.

## Introduction

The phylum Apicomplexa comprises a vast group of single-celled protozoa that pose a huge impact on global health and the economy by causing diseases such as malaria, toxoplasmosis, cryptosporidiosis, and coccidiosis. *Plasmodium*, the causative agent of malaria, alone infects 247 million people and claims 600,000 lives annually. A distinctive feature of most <u>apico</u>mplexan parasites is the presence of a non-photosynthetic <u>plast</u>id called the <u>apicoplast</u>, which originated from secondary endosymbiosis and is therefore surrounded by four membranes^1^. The apicoplast in malaria parasites is essential for parasite survival and replication throughout its complex lifecycle. Notably, antibiotics such as tetracycline^2^ were used to treat malaria patients decades before the apicoplast was even identified^3,4^.

Due to its endosymbiotic origin, the apicoplast features many bacterial or plant-like pathways that synthesize critical metabolites, including fatty acids, lipoic acid, isoprenoid precursors, iron sulfur clusters, and heme^5^. During the asexual blood stage, where malaria clinical symptoms arise, the apicoplast undergoes a growth and fission cycle to divide into 16-32 copies to ensure that each progeny inherits a functional apicoplast. Despite being essential in the asexual blood stage, the apicoplast appears to be undergoing reductive evolution in a “*going, going, gone”* manner, as many of its synthetic pathways—such as FASII and heme biosynthesis—are dispensable in this stage^6^. Strong support for this idea came from a 2011 study^7^ that demonstrated asexual *P. falciparum* could survive without an apicoplast when treated with antibiotics and rescued by sufficient amounts of isopentenyl pyrophosphate (IPP, 200 µM), the product of the non-mevalonate/methylerythritol phosphate (MEP) pathway present within the apicoplast. IPP, a five-carbon isoprene, is used for synthesizing ubiquinone and long-chain isoprenoids involved in protein prenylation and glycosylation (via dolichols). The ability of parasites to survive indefinitely under antibiotic treatment when supplemented with IPP suggested that the sole essential function of the apicoplast in asexual blood stages is to synthesize a single molecule: IPP^7^. These IPP-rescued “apicoplast-minus” parasites lack the 35 kb apicoplast genome and an intact organelle but possess numerous vesicles, which were proposed to be transport vesicles^7^.

This “apicoplast minus” theory was groundbreaking, as it implied that, given access to IPP from alternative sources, malaria parasites could evolve to lose the entire organelle at least in one stage^6^, mirroring the situation observed in *Cryptosporidium*, an apicomplexan parasite that has truly lost its apicoplast. Notably, *Plasmodium* and *Cryptosporidium* are estimated to have diverged approximately 420 million years ago^8^. Consequently, the 2011 study suggested that a major evolutionary event, typically taking millions of years, could occur within a laboratory setting within just a matter of a few days. Since then, the concept of “apicoplast-minus” *P. falciparum* has been widely accepted.

However, here we show that the apicoplast has never truly disappeared in the “apicoplast minus” *P. falciparum* parasites, challenging the long-standing belief that the organelle can be entirely bypassed by IPP supplementation.

## Results

In the asexual *P. falciparum*, IPP rescue can be achieved by two ways: direction addition (200 µM) or synthesis from a precursor (mevalonate, 50 µM) in an engineered parasite line PfMev^9^. Compared to direct IPP supplementation (approximately $3500 per liter of medium, Sigma-Aldrich), PfMev reduces the cost by several orders of the magnitude (approximately $0.06 per liter of medium). Both methods are equally effective in inducing the “apicoplast minus” phenotype. Additionally, PfMev is well-suited for live-cell imaging, as its apicoplast is labeled by sfGFP (super folder GFP).

PfMev was treated with erythromycin and mevalonate to induce the “apicoplast minus” phenotype and continuously cultured for several months. In the resulting “apicoplast minus” PfMev, referred to hereafter as PfMev(-), we confirmed the absence of the apicoplast genome, the presence of numerous green vesicles, and no cross-contamination of any apicoplast-containing parasites, as PfMev(-) quickly died after mevalonate withdraw (data not shown). We stained PfMev(-) with MitoTracker and performed live-cell microscopy at different time points over the 48-hour asexual lifecycle. Our observations challenged the prevailing theory of “apicoplast minus” (**Figure 1**). According to this theory^7,10^, ring-stage parasites should contain no apicoplast due to the organelle’s loss in the previous cycle. In contrast, we observed a single green structure in ring-stage parasites. During the trophozoite-stage, we detected numerous apicoplast-derived vesicles and the absence of an intact apicoplast, closely resembling the typical “apicoplast-minus” phenotype as observed in 2011^7^. Notably, these vesicles appeared to be randomly distributed throughout the parasite cytosol. However, in the schizont-stage, the apicoplast-derived vesicles started to “colocalize” with the mitochondrion, suggesting non-random distribution. The close association of the vesicles with the segmented mitochondria became even more pronounced in the late schizont-stage. Upon egress, both structures were packed into merozoites prior to invasion.

**Figure 1.**
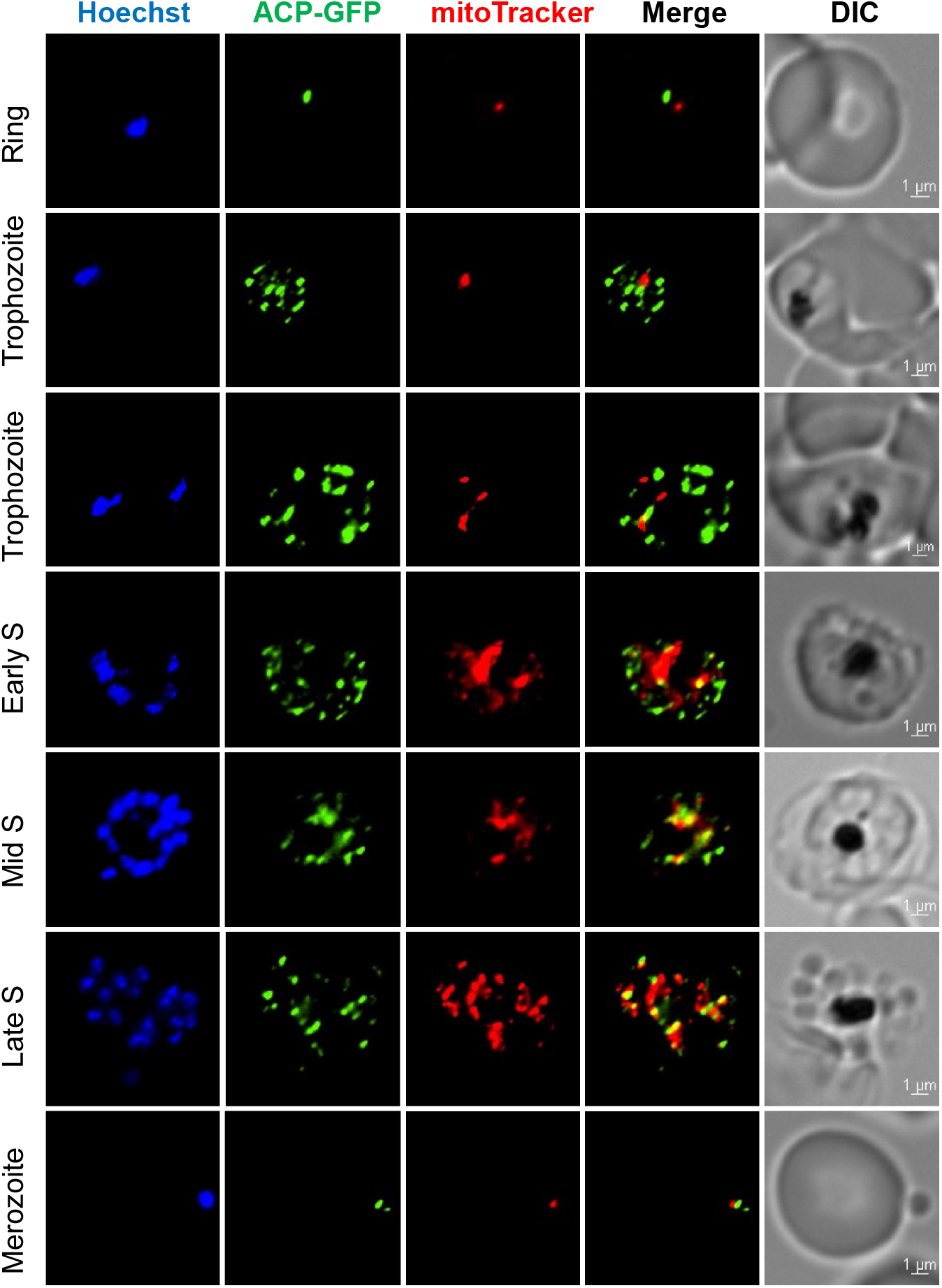
The disrupted apicoplast in the “apicoplast minus” *P. falciparum* is inherited. Live-cell imaging monitors the disrupted apicoplast in PfMev(-). Hoechst, DNA; ACP-sfGFP, apicoplast; MitoTracker, mitochondrion. Pearson’s correlation coefficient of parasites (n=10) at each stage: Ring (0.440±0.378), Trophozoite (0.560±0.127), Early S (early schizont, 0.776±0.034), Mid S (middle schizont, 0.850±0.025), LS (late schizont, 0.945±0.008), Merozoite (0.915±0.018). Bars, 1 µm. This experiment was repeated four times.

These data demonstrate that the disrupted apicoplast in the “apicoplast minus” parasites is inherited.

To further verify the nature of the apicoplast-derived vesicles at higher resolution, we performed immuno-EM on PfMev(-) using an antibody against ACP, acyl carrier protein (the apicoplast marker). As shown in **Figure 2**, we detected two distinct types of ACP-positive structures in the trophozoite-stage parasites. The smaller structures, approximately 50-100 nm in diameter, are circular and appear to be enclosed by a single membrane. These vesicles likely contain nuclear-encoded proteins, responsible for delivering cargo to the apicoplast. Strikingly, the larger structures (>200 nm) are enclosed by four membranes, unequivocally identifying them as the disrupted apicoplasts rather than unknown entities. The presence of four-membrane-bound structures strongly suggests that the apicoplast never truly disappears in the antibiotics treated, IPP-rescued parasites. Instead, it appears to undergo an unknown transformation, becoming smaller in size and exhibiting multiple copies per parasite. We have named this genome-less apicoplast derivative the apicosome, drawing parallels with the genome-less mitochondrial derivative, the mitosome, found in deeply divergent eukaryotes such as *Entamoeba histolytica*.

**Figure 2.**
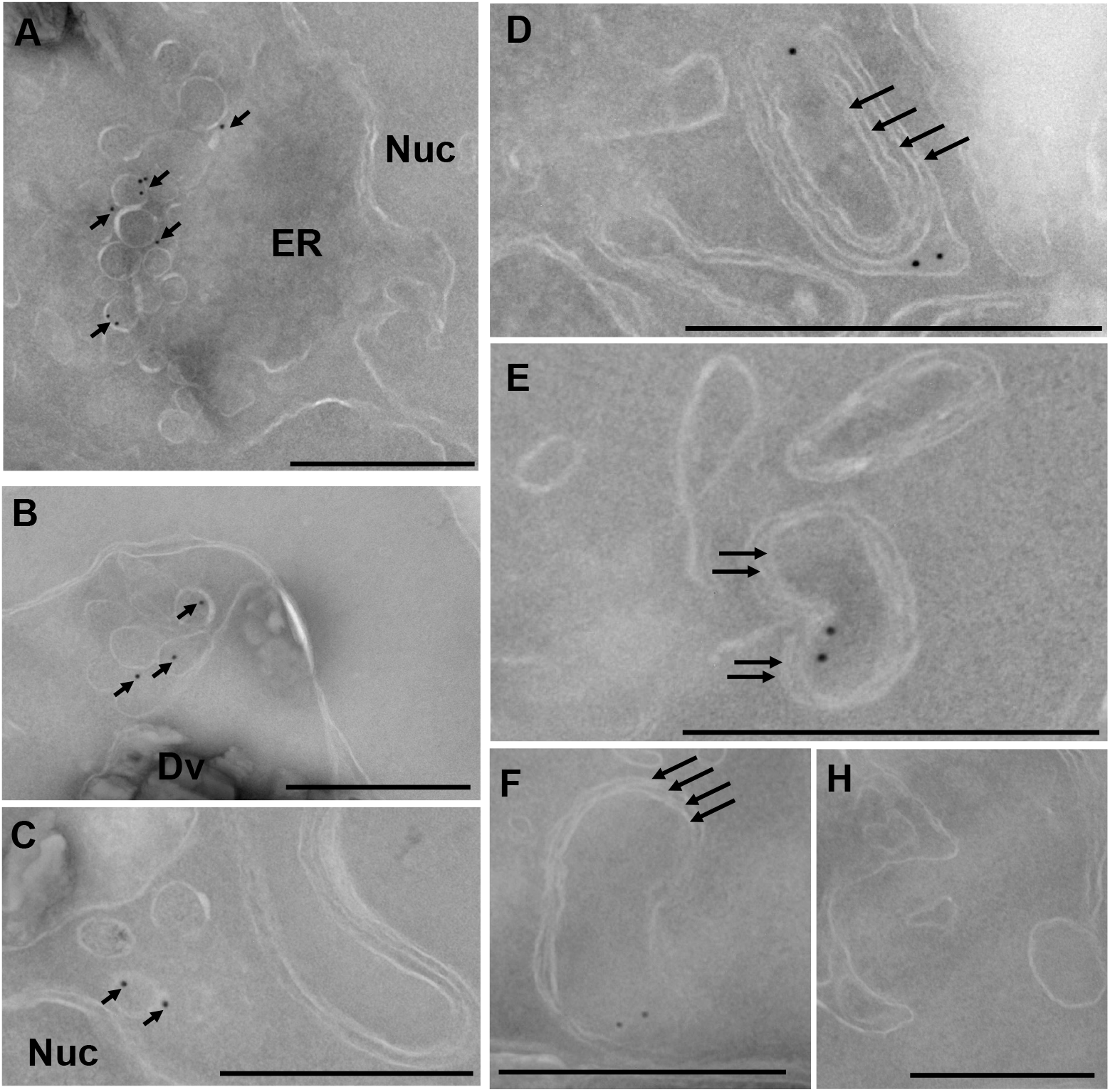
Detection of structures surrounded by four membranes in the “apicoplast minus” parasites by immuno-EM. (A-C), ACP positive structures surrounded by a single membrane (short black arrows). Nuc, nucleus; ER, endoplasmic reticulum; Dv, digestive vacuole. (D-F), ACP positive structures surrounded by four membranes (long black arrows). ACP, acyl carrier protein. H, negative control with no primary antibody. Bars = 500 nm. This experiment was repeated twice.

## Discussion

Our data suggest that the apicoplast is not entirely lost but instead persists in a distinct, diminished form when the parasites encounter antibiotic treatment and IPP rescues. We propose that apicosomes continue to serve essential biochemical and/or structural functions, acting as barriers to the complete loss of the apicoplast in the asexual malaria parasites.

We think the apicosome likely contains core essential proteins that remain indispensable even after apicoplast disruption. Indeed, one such protein has already been discovered: DPCK (dephospho-CoA kinase), the final enzyme in the CoA biosynthesis pathway. Deletion of the apicoplast-located DPCK is lethal in PfMev even with IPP supplementation^11^ as CoA is used by multiple subcellular compartments, including the endoplasmic reticulum (for fatty acid elongation) and the nucleus (for histone acetylation). Further studies are needed to determine the apicosome’s proteome to identify additional essential functions that cannot be bypassed by IPP supplementation. Beyond biochemical roles, we speculate that apicosomes may play crucial roles in organellar segregation during parasite division. A recent study suggests that the apicoplast is more closely associated with the parasite nucleus (within 50 nm) than with the mitochondrion (typically > 500 nm)^12^, indicating that the apicoplast, not the mitochondrion, maybe responsible for leading organellar segregation. If true, this could explain why the apicoplast can never be truly lost because its absence would disrupt mitochondrial distribution and cause parasite death. Clearly, further investigation is needed to unravel the role of apicosomes in parasite division.

In summary, our discovery of apicosomes in IPP-rescued *P. falciparum* highlights the significance of the apicoplast in malaria parasites and underscores the complexity of apicoplast function and maintenance.

## Materials and Methods

Methods of parasite culture, synchronization, and immuno-EM have been described previously^13,14^. Live cell imaging was performed with a Nikon Ti microscope and images were processed with the Nikon NIS elements software.

## ACKNOWLEGEMENTS

We are grateful to Dr. Wandy Betty (Washington University in St Louis) for performing immuno-EM studies and Dr. Akhil B. Vaidya for critical discussion. We also thank ChatGPT for correcting minor grammatic issues. This study is supported by NIH grants to Dr. Hangjun Ke (R21 AI156735, R01 AI184855) and Dr. Sean Prigge (R01AI125534).

